# Automated In Vitro Wound Healing Assay

**DOI:** 10.1101/2023.12.23.573213

**Authors:** Jillian Cwycyshyn, Cooper Stansbury, Walter Meixner, James B. Hoying, Lindsey A. Muir, Indika Rajapakse

## Abstract

Restoring the epidermal barrier after injury requires spatial and temporal orchestration of migration, proliferation, and signaling across many cell types. The mechanisms that coordinate this complex process are incompletely understood. In vitro wound assays are common model systems for examining these mechanisms in wound healing. In the scratch assay, a cell-free gap is created by mechanical removal of cells from a monolayer, followed by monitoring cell migration into the gap over time. While simple and low-cost, manual scratch assays are limited by low reproducibility and low throughput. Here, we have designed a robotics-assisted automated wound healing (AWH) assay that increases reproducibility and throughput while integrating automated live-cell imaging and analysis. Wounds are designed as computer-aided design (CAD) models and recreated in confluent cell layers by the BioAssemblyBot (BAB) 3D-bioprinting platform. The dynamics of migration and proliferation in individual cells are evaluated over the course of wound closure using live-cell fluorescence microscopy and our high-performance image processing pipeline. The AWH assay outperforms the standard scratch assay with enhanced consistency in wound geometry. Our ability to create diverse wound shapes in any multi-well plate with the BAB not only allows for multiple experimental conditions to be analyzed in parallel but also offers versatility in the design of wound healing experiments. Our method emerges as a valuable tool for the automated completion and analysis of high-throughput, reproducible, and adaptable in vitro wound healing assays.

## Introduction

Cutaneous wound healing requires coordinated signaling across cell types in four major phases: hemostasis, inflammation, re-epithelialization, and tissue remodeling [1–4] (Figure 1). Injury to the skin triggers the formation of a fibrin clot to initially reestablish the epidermal barrier. Immune cells attracted to the site of injury release bioactive factors that stimulate migration and proliferation of fibroblasts and keratinocytes. Activated fibroblasts reconstruct the extracellular matrix (ECM), while keratinocytes re-epithelialize the tissue [4, 5]. Disruptions to this system can lead to excessive scar formation or the development of diabetic ulcers, keloids, and other chronic non-healing wounds [4, 6, 7]. The repercussions of impaired wound healing also extend beyond chronic wounds to severe skin injuries where precise cellular responses are paramount for optimal tissue regeneration. In cases of deep skin trauma and thermal burn wounds, the intricacies of the wound healing cascade become even more crucial.

**Figure 1:**
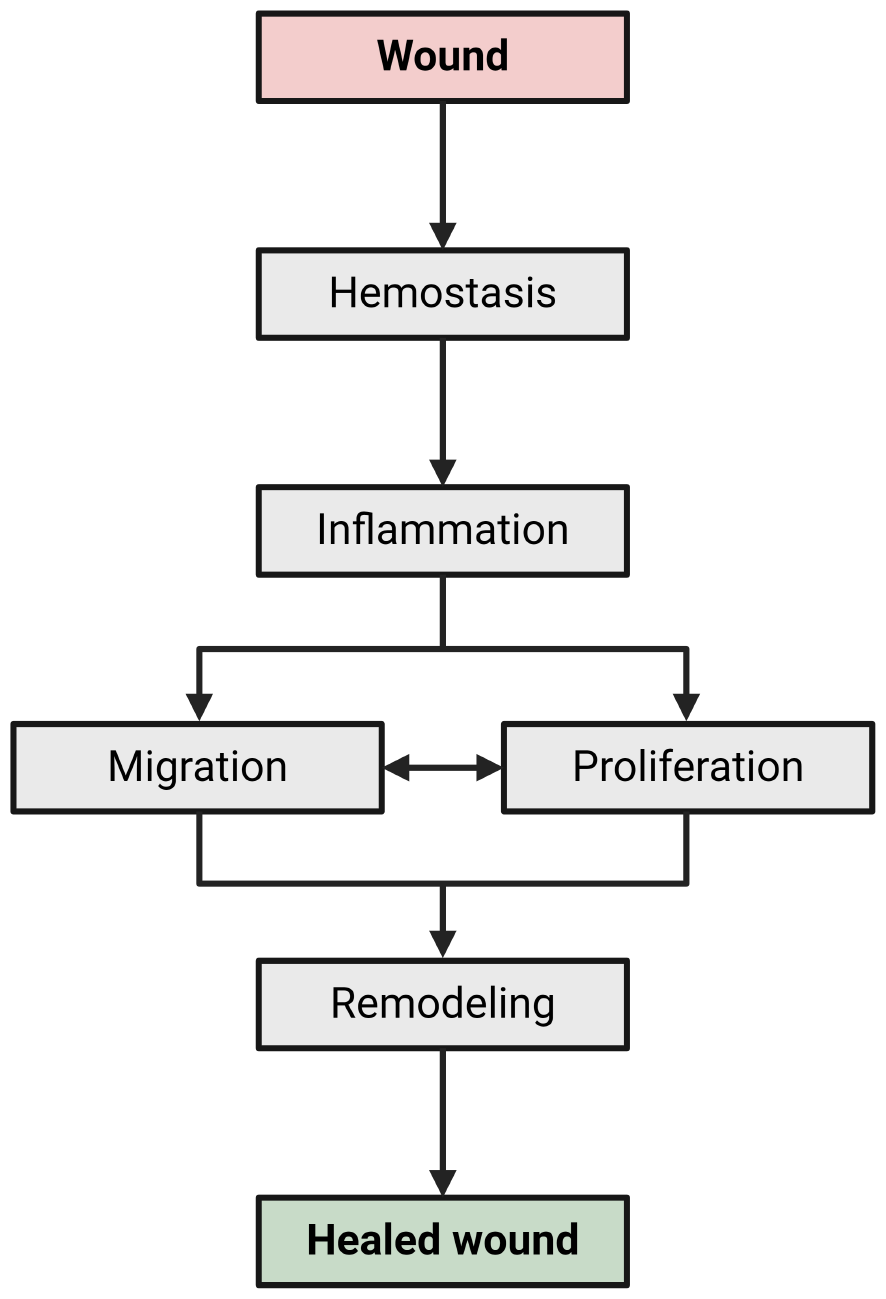
Overview of the wound healing process. Hemostasis, inflammation, re-epithelialization, and tissue remodeling are overlapping phases that take place throughout the course of wound healing. During re-epithelialization, cellular migration and proliferation are delicately balanced to close the wound and restore tissue function.

Two-dimensional in vitro wound healing assays incorporating primary skin cells are frequently used to investigate migration and proliferation dynamics. Traditional methods, like the scratch assay, are simple and cost-effective but suffer from limited reproducibility, low throughput, and inflexibility in experimental design [8]. Likewise, while culture insert or barrier-based methods have high reproducibility, they are low throughput and must be frequently replaced [8, 9]. Several technologies have been developed to automate the creation of scratch wounds in 96-well plates [10, 11]. However, these methods create uniform scratches in all wells simultaneously which limits versatility in testing multiple experimental conditions or diverse wound types (e.g. incisions, lacerations, punctures).

The integration of robotic systems in biological research has revolutionized our ability to conduct complex assays [12, 13]. Within the context of wound healing, robotics-assisted automation provides a compelling opportunity to develop a reproducible, high-throughput, and adaptable wound healing assay. Herein, we present the automated wound healing assay (AWH)—a robotically-controlled approach to consistently create monolayer wounds and analyze cellular migration and proliferation across those wounds throughout the course of wound closure. Our method enables researchers to implement a wound healing assay in any-sized cell culture well plate, with the novel ability to generate customized wounds using an intelligent six-axis robotic arm, termed the BioAssemblyBot (BAB). The BAB is designed for 3D bioprinting and the completion of complex assays. Using the BAB’s built-in printing workflow and 3D modelling software, wounds are designed as 3D shapes flattened onto the culture surface of a pre-calibrated well plate. CAD wound models are then sent to the BAB as a “3D print”, where the AWH is executed and the designed wounds are created in confluent cell layers. Wound closure is monitored over time (typically 2-4 days) via live-cell fluorescence microscopy, enabling both the migratory and proliferative activities of individual cells to be tracked throughout the course of healing.

## Results

### Automated wound healing assay design

For AWH, we programmed BAB to mechanically remove cells from a monolayer with consistent speed and pressure using a dispensing tip on its Printing Tool (Figure 2A). This approach produced highly consistent wound dimensions and minimal damage to the cells and culture surface.

**Figure 2:**
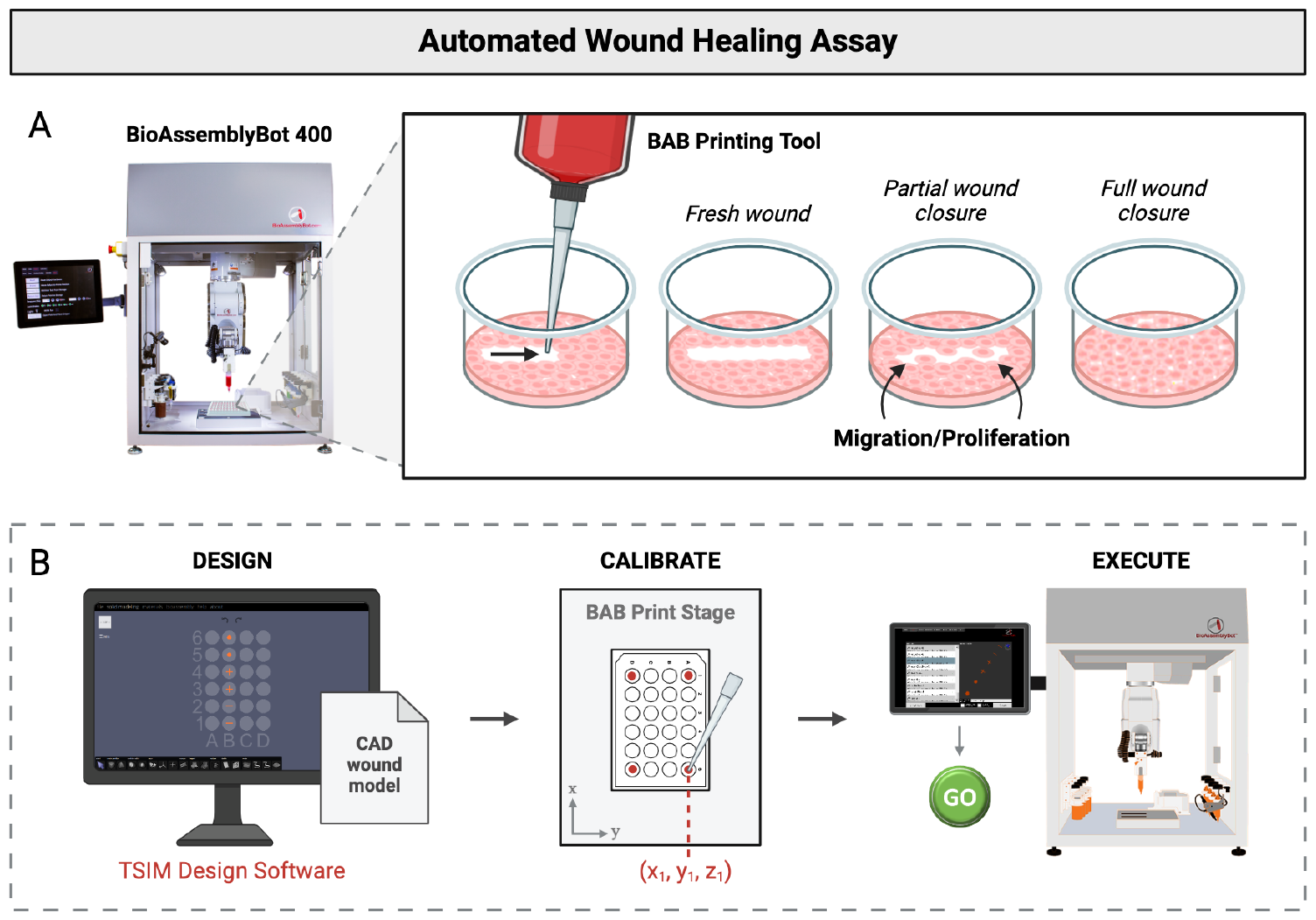
Overview of automated wound healing. The BioAssembly Platform, including the BioAssemblyBot 400 (BAB) and the TSIM design software, was used for the automation of wound healing assays to increase reproducibility, scalability, and controllability of wound healing experiments. **(A)** BAB is programmed to mechanically remove userdefined regions from cell monolayers. Processes such as migration and proliferation can be monitored during wound closure. **(B)** Workflow to implement AWH. First, 3D models of the wounds are designed using TSIM software. Next, the well plate is calibrated with BAB. This involves determining the x, y, and z coordinates of each corner of the plate relative to the print stage with BAB’s Printing Tool. Lastly, the CAD wound files are exported to BAB, where AWH is executed in the BAB user interface.

AWH was implemented in three parts: 1) wound design, 2) plate calibration, and 3) execution (Figure 2B). Wounds were first designed as 3D CAD models, where wound shape, size, and placement within the wells were customized. Plate calibration was performed on the same type of multi-well plate as the prepared cultures and the same type of dispensing tip on the BAB arm. X, y, and z coordinates were programmed for the center of all wells relative to the print stage. The 3D wound models were then sent to the BAB, where the calibrated plate was selected as the container for printing, and the AWH assay was executed.

### Automated wound healing generates consistent wound dimensions with control over wound shape

To compare the performance of the AWH assay against the standard scratch assay, simple scratches were applied to human fibroblasts by the BAB and manually. Images of each well were taken at 5x magnification, and manual- and BAB-generated scratches were compared for their consistency and positioning. The AWH assay produces wounds with increased consistency in wound shape across all wells compared to wounds that were created manually (Figure 3A). We observed smaller deviations in scratch width for scratches generated by the BAB (*σ* = 44.9 *µ*m) compared to manually-generated scratches (*σ* = 103.1 *µ*m) (Figure 3B). The positioning of the scratches across all wells was more variable for manual scratches compared to the AWH assay, which exhibited uniform positioning of each scratch near the center of the well (Figure 3C). The BAB’s 3D-printing workflow can create complex wound shapes with high precision (Figure 3D). To evaluate the impact of geometry on wound closure, we designed an AWH assay to create different shapes and performed live-cell imaging at 20 minute intervals over 88 hours. Representative images of the circle and triangle wounds closing over time are depicted in Figure 3E. As expected, the area of the wound determined wound closure time (Table 1). This emphasizes the importance of reproducibility in wound healing assays—manually-generated wounds with higher error in wound geometry can affect the experimental outcomes. Thus, in a given experiment, wounds need to be carefully designed and created in order to make accurate comparisons within and across different conditions.

**Table 1:**
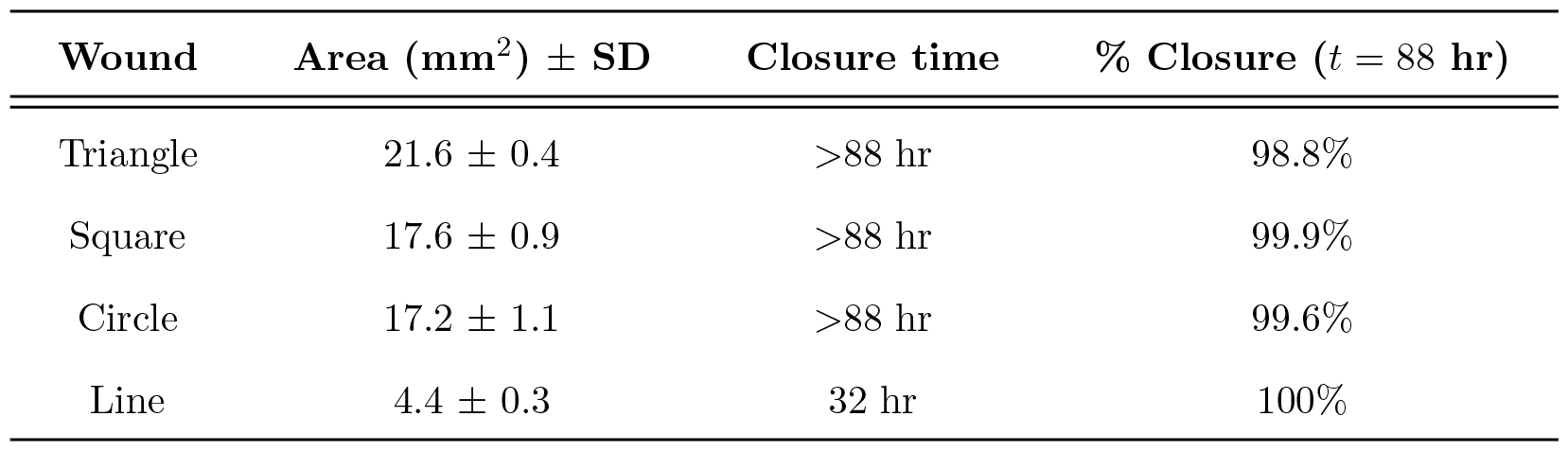
Average area and wound closure times for different wound shapes (n = 3).

**Figure 3:**
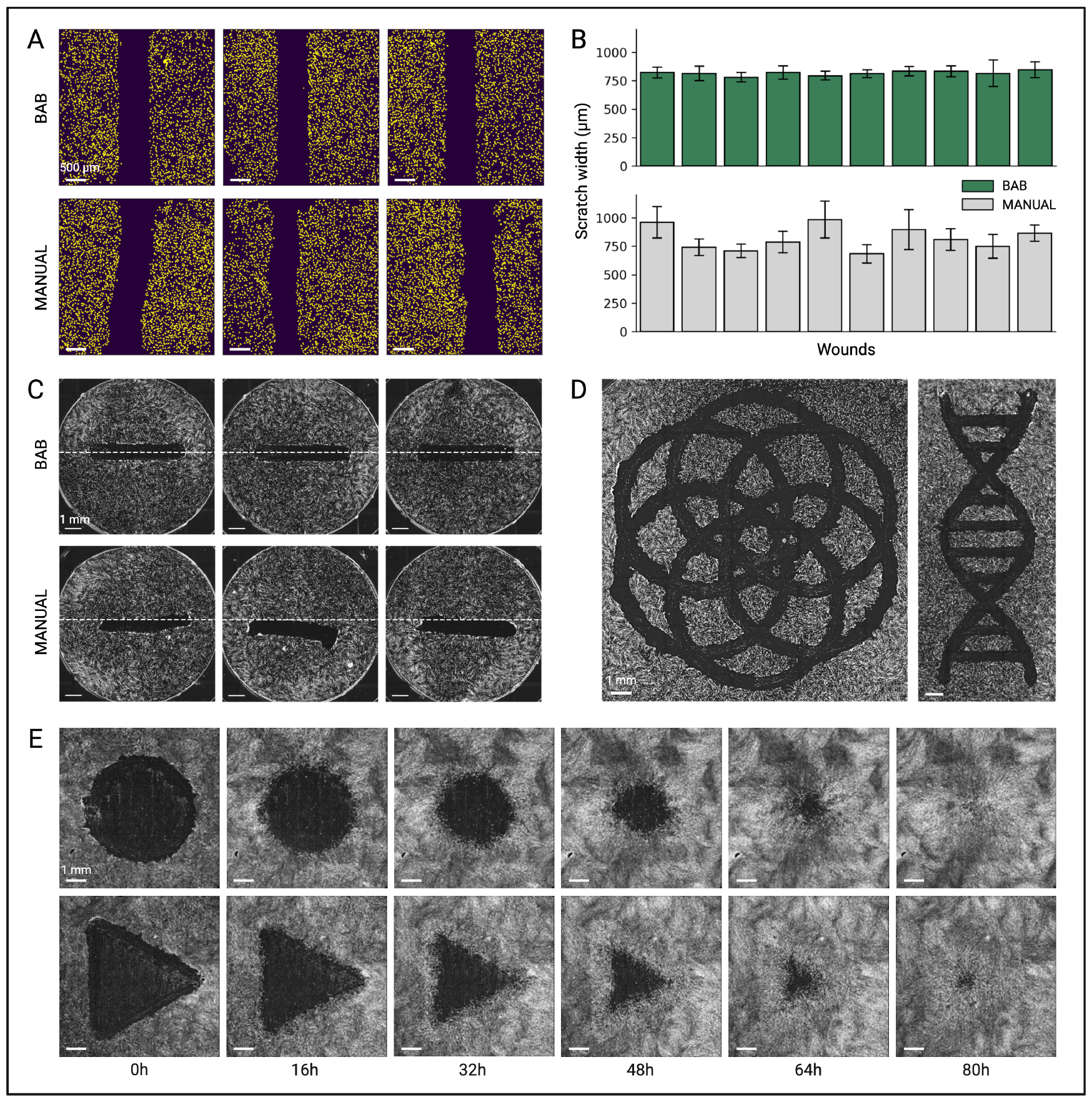
The AWH generates more consistent wounds compared to the standard scratch assay. **(A)** Representative images of simple scratches made by the BAB (top) and by hand (bottom). BAB-generated scratches exhibit more consistency compared to manually generated wounds. Scale bar = 500 *µ*m. **(B)** Scratch width measurements. Scratch widths and standard deviations were calculated as the distance between each wound edge along the full length of the scratch. Each bar represents one wound. **(C)** Representative full well images of scratches made by the BAB (top) and by hand (bottom). White dashed lines indicate the center of the well. Scale bar = 1 mm. **(D)** Representative images of complex wound shapes created by the BAB. Scale bar = 1 mm. **(E)** Image montage of wound closure over time for circle and triangle wound shapes. Scale bar = 1 mm.

### Live-cell fluorescent microscopy enables monitoring of migration and proliferation in individually tracked cells

Live-cell imaging enables capture of proliferation and migration dynamics after wounding. We leveraged BAB’s built-in capacity for interfacing with other lab equipment to couple AWH with live-cell imaging to build a fully automated workflow (Figure 4A).

**Figure 4:**
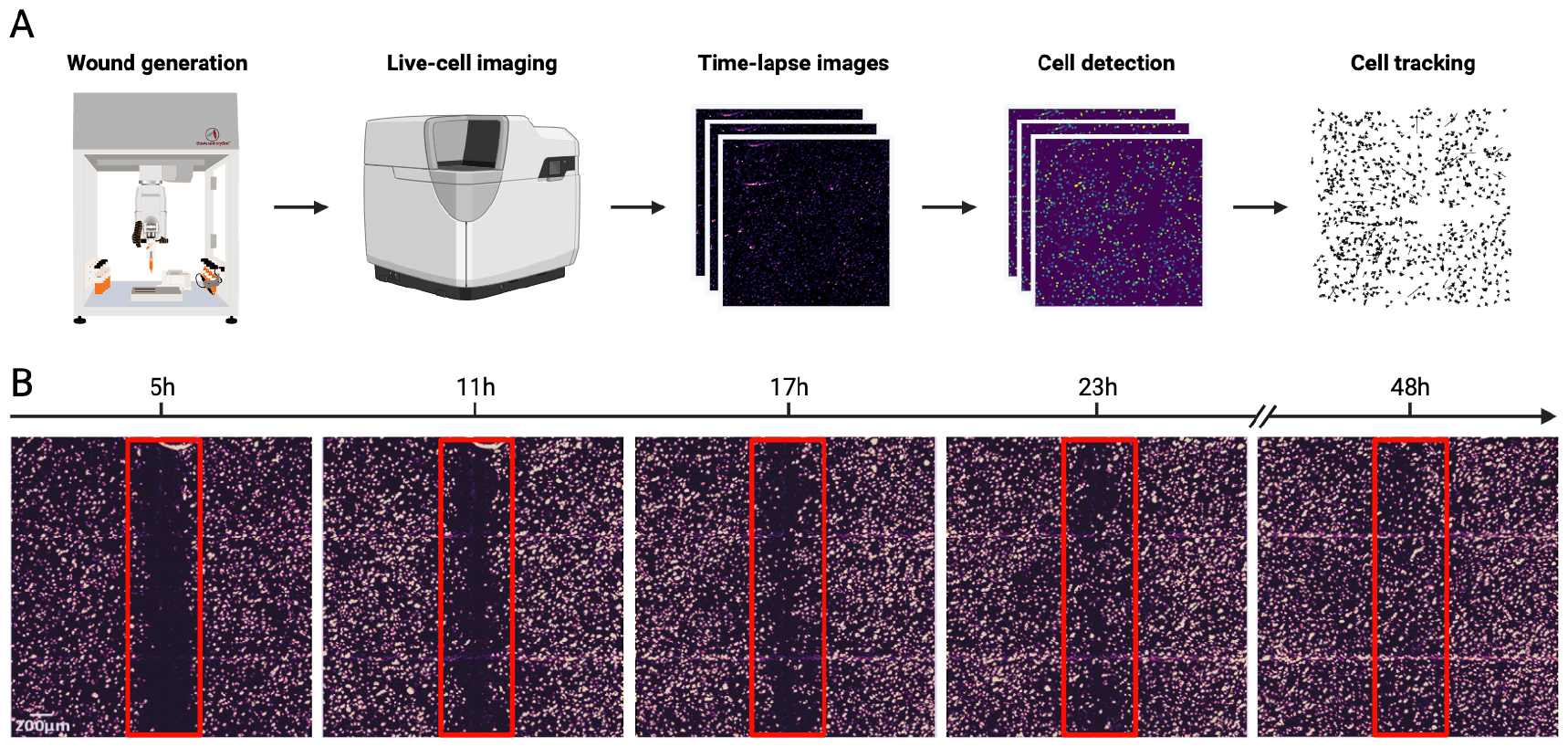
Workflow for the automated wound healing assay and time-lapse image analysis. **(A)** The AWH is performed as described in Figure 2. By use of the BioApps Maker workflow automater, the BAB is programmed to transfer the well plate to the Zeiss Celldiscoverer7 (CD7) and begin image acquisition. Images are taken at 20-30 minute intervals over the course of wound closure. Individual cells are detected in each image, and the migration trajectory for each cell is tracked over time. **(B)** Representative images of wound closure over time for a simple scratch wound. Scale bar = 200 *µ*m.

As a proof of concept, AWH was performed on human fibroblasts. We programmed the BAB to transfer the “wounded” well plate to the Zeiss Celldiscoverer 7 (CD7) live-cell fluorescent microscope with its Pick n’ Place tool. Time-lapse images of the cells were captured at 5x magnification every 30 minutes over 48 hours. The acquired images were run through our image processing pipeline, which detects and tracks individual cell nuclei over time (Figure 4B).

## Discussion

The automated wound healing assay demonstrates notable advantages in terms of reproducibility, scalability, and experimental flexibility compared to traditional methods. The robotic control of the assay not only minimizes human variability but also ensures consistent wound creation and monitoring. The scalability and flexibility of our approach allows for high-throughput experimentation, making it adaptable to various experimental conditions and wound types. The incorporation of live-cell fluorescence microscopy in our automated workflow opens up avenues for assessing additional biological processes during wound healing. This real-time imaging capability enables the observation of dynamic cellular events, such as cell cycle phase transitions [14] or epithelial-to-mesenchymal transition (EMT) dynamics [15], providing a more comprehensive view of cellular behaviors in response to injury.

An exciting prospect of our automated workflow is its applicability to cellular reprogramming studies geared towards accelerating the wound healing process. In previous work, we have successfully reprogrammed human fibroblasts to embryonic stem cells, muscle cells, and other cell types utilizing our in-house data-guided control algorithm [16]. This framework employs a multi-way dynamical systems approach to predict combinations of TFs, as well as the specific time point in which TF activation or suppression will have the greatest effect, for converting one cell type into another desired phenotype. By leveraging the precision and control offered by the AWH assay, we envision that future iterations of our model can be refined to explore the modulation of cellular states during wound healing through transcription factor-guided therapy (Figure 5).

**Figure 5:**
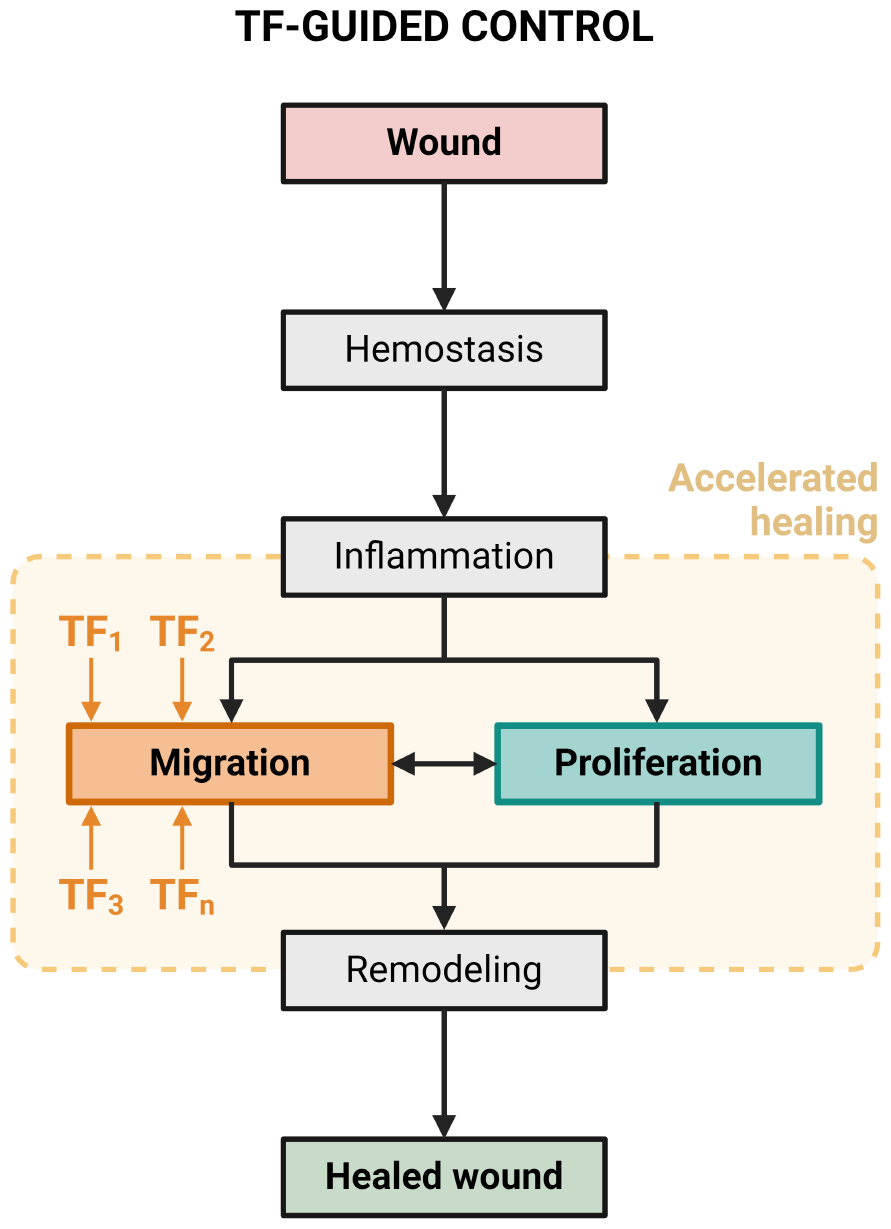
The wound healing process under transcription factor-guided control. We envision that we can expedite the wound healing process through a data-guided cellular reprogramming approach. Delivering algorithmicallypredicted transcription factors to wounded cells has the potential to enhance cellular migration and proliferation, thus promoting accelerated healing. proximal to the injury site lose their epithelial characteristics and adopt a motile mesenchymal-like program, while epidermal cells distal to the wound edge begin to proliferate. As the underlying tissue repairs, keratinocytes repopulate the wound bed and reestablish their epidermal signature [5, 18].

Wound healing has long been recognized as a process akin to cancer metastasis [17, 18]. Keratinocytes have the innate ability to partially and reversibly transition into another phenotype. This feature of wound healing resembles that of the epithelial-mesenchymal transition (EMT), a process in which an epithelial cell acquires a mesenchymal phenotype. In cancers, EMT is known to be a key player in tumor progression, metastasis, and chemo-therapeutic resistance [19, 20]. In wound healing, this partial EMT is crucial to the restoration of epidermal barrier function. Keratinocyte cells Although it is known that chromosomal aberrations and certain core EMT genes are not apparent in physiological re-epithelialization [18], the evident similarities between cancers and wound healing mechanisms prompts the notion that deepening our understanding of the intricate balance between migration and proliferation during wound healing could augment our abilities to understand cancer metastasis. Human-relevant in vitro models have recently been approved by the FDA as a viable alternative to animal testing in preclinical drug evaluations [21]. Recognizing the well-documented pharmacogenomic variations between animal models and humans, the use of biomimetic cell-based assays for drug safety and efficacy testing represents a significant step toward more accurate predictions of human therapeutic responses. In the context of wound healing and cancers, our automated method can be harnessed to screen for potential EMT regulators and, in the long term, to evaluate the efficacy of anti-cancer drugs targeting this process.

## Conclusion

We have developed an automated workflow for the execution and analysis of in vitro wound healing experiments. Our automated wound healing assay not only addresses the limitations of traditional methods but also opens up avenues for exploring diverse biological phenomena during wound healing and advancing research in regenerative medicine and cancer biology. The reproducibility, scalability, and adaptability of our approach, coupled with its potential applications, position it as a valuable tool in the broader landscape of experimental techniques.

## Methods

### Automated wound healing assay

The steps described below were completed for each run of the AWH assay reported in the Results. Multi-well plates from 12 - 96 wells were tested in the AWH assay. The plate calibration steps were completed once for each plate size used, while the tip to stage offset was determined for each new dispensing tip.

### Wound design

Tissue Structure and Information Modeling (TSIM) software v1.1.227 (Advanced Solutions Life Sciences, LLC) was used for design of the wounds. The plate type was selected and settings included “print continuously”, “flatten”, and the printer default for “move between layers.” In a new sketch, the wound shapes were drawn using the software’s sketching tools. Finalized wound shapes were then copied into other wells as desired for replicates. The z-depth of all objects was set to 0.0001. In the material settings, printing pressure was set to 0 psi, and printing speed and acceleration were set to 3 *mm/sec* and 10 *mm/sec*^2^, respectively. The design file was saved and sent as a print job to the BAB human machine interface (HMI).

### Plate calibration

For plate calibration, multi-well plates were placed on the BAB Print Stage and de-lidded. In the BAB HMI, the BAB Printing Tool was retrieved and a new container for printing was created for each type of plate. The number of rows and columns were input, and the Printing Tool tip was manually positioned in the center of the first well, designated A1, at an arbitrary z-coordinate since z coordinates were determined later during tip to stage offset calibration. The x, y, and z-coordinates were recorded for A1 and the process was repeated for the remaining three corner wells. The coordinates were then calculated and stored for all wells using the HMI settings Update All Wells (Relative) and Calculate Wells from Extents. Calibration for tip to stage offset ensured contact between the dispensing tip of the BAB Printing Tool and the plate surface. The BAB Printing Tool with a tip was manually positioned to have full contact with the surface of the plate, ensuring no deformation of the tip. The measured tip offset was recorded for use during print execution.

### Execution

The assay was executed using the Print interface of the BAB HMI. The measured tip offset was loaded in the calibration settings alongside the design file received from the TSIM application. The calibrated well plate was selected as the container for printing and the location of the BAB Printing Tool was given.

### Cell culture

Human BJ fibroblasts (ATCC CRL-2522) were cultured on standard cultureware in Dulbecco’s Modified Eagle’s Medium (DMEM, Gibco 11965-092) with 10% Fetal Bovine Serum (FBS, Corning 35-015- CV), 1% MEM Non-Essential Amino Acids (NEAA, Gibco 11140-050), and 1% penicillin-streptomycin (P/S, Gibco 15140122). Cells were incubated at 37°C in 5% CO_2_, and media were exchanged every 48 hours. For the AWH assay, BJ Fibroblasts were seeded at densities ranging from 0.22 - 0.63 x 10^5^ cells/*cm*^2^ in 12-, 24-, 48-, or 96-well plates. After 24 hours, cells were incubated in normal media containing 0.02 *µ*M Hoechst 33342 (Enzo, ENZ-52401) for 2 hours, followed by executing the AWH assay. Wells were then washed with PBS. FluoroBrite DMEM (Gibco, A18967-01) with 10% FBS, 1% NEAA, and 1% P/S was added to all wells prior to image acquisition.

### Image acquisition

The Ziess Celldiscoverer 7 (CD7) live-cell imaging system was used to automate capture of time-lapse images during wound closure. Oblique contrast and fluorescence microscopy was performed with a Plan-Apochromat 5x/0.35 objective and 0.5x or 1x tube lens. Images were taken using an Axiocam 506 with 14 bit resolution. Cells were imaged at 37°C in 5% CO_2_. Images were captured every 20 or 30 minutes over the duration of wound closure. For each wound, a multi-channel time-series ome.tiff file was prepared in the Zen Blue 3.0 software and exported for downstream analysis.

## Image processing

### Image analysis

Raw images were preprocessed using `ImageJ` and `Python`, and analyses were performed with `MATLAB` and `Python`. All scripts may be found at the following URL https://github.com/jrcwycy/wound_healing.

For wound area calculations, raw oblique images were first preprocessed in ImageJ. Wounds were identified using the `Image Segmenter` application in `MATLAB` and exported as binary masks. Wound areas were then quantified over time for each shape. For scratch width analysis, individual cell nuclei were segmented from raw H3342 images using `StarDist` [22], a deep-learning based method for object detection and segmentation. Isolated segmentations were then filtered out of the wound bed images using a nearest neighbors approach [23]. To identify wound edges, contours that separated areas of high cell density from areas of low cell density were detected using `scikit-image` [24]. The line of best fit for each contour served as the wound edge for subsequent analyses. Scratch widths were calculated as the distance between the two wound edges along the entire length of the scratch for each image.

### Automated cell tracking

To automate analysis of wound healing experiments, we constructed an image processing pipeline using the `Python` framework `Snakemake` [25, 26]. The pipeline is designed to manage parallel processing of large time-series imaging data in a high-performance computing environment. Briefly, the pipeline produces nuclear segmentations at each timestep using `StarDist` [22] and predicts cellular movement using a Bayesian single cell tracking approach [27]. Inputs to the pipeline are multi-channel time-series `ome.tiff` files and a set of user-defined parameters controlling the behavior of different filtering and analysis operations [24]. The outputs of the pipeline are properties of cell nuclei at each time step, and nuclear linkages between time-steps which we refer to as ‘tracks.’ We describe the operations of the pipeline in Algorithm 1 on a single input. Note that the pipeline may be run on a set of input images. All software for automated image prepossessing and analysis may be found at the following URL https://github.com/CooperStansbury/pip-fucci.

## Supporting information

Supplemental Images and Data

## Supplementary Material

See the supplementary material for images and data that support the findings of this study.

### Algorithm 1 Automated Cell Tracking

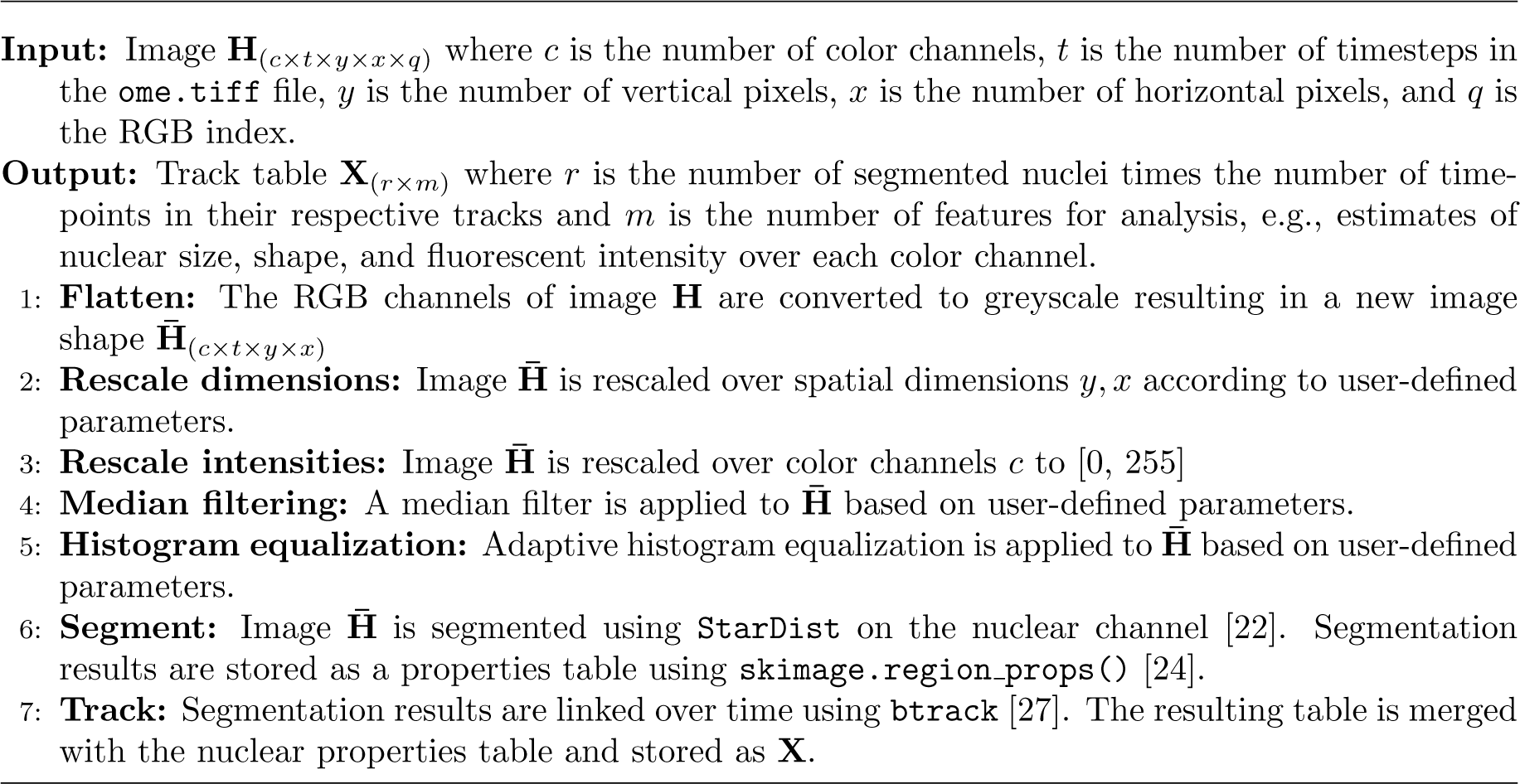

## Acknowledgments

We thank Gilbert S. Omenn for critical reading of the manuscript. This work is supported by the Air Force Office of Scientific Research under award number FA9550-22-1-0215, Defense University Research Instrumentation Program (DURIP) 2018, DURIP 2022, and the National Science Foundation (NSF) under award number 2225568. Figures were created with BioRender.com.

## Author Declarations

### Ethics Approval

Ethics approval is not required.

### Author Contributions

J.C., J.H., L.M., and I.R. designed research; J.C. and W.M. performed research; J.C., C.S., J.H., and I.R. contributed new reagents/analytic tools; J.C., C.S., W.M., and L.M. analyzed data; J.C., C.S., J.H., L.M., and I.R. wrote the paper.

### Data Availability

The data that support the findings of this study are available within the article and its supplementary material. Additional supporting images are available from the corresponding authors upon reasonable request.

